# Canopy cover and structural complexity affect the phylogenetic composition of a pond-dwelling Neotropical anuran metacommunity

**DOI:** 10.1101/252478

**Authors:** Diogo B. Provete

## Abstract

Phylogenetic information has been increasingly included into (meta)community assembly studies. However, recent studies have challenged the framework commonly used to infer processes from phylogenetic structure. Amphibians are good model organisms to study processes promoting structure in metacommunities, since they are subjected to different environmental and spatial processes throughout their biphasic life cycle. Pond canopy cover is one of these environmental factors that strongly influence the distribution of species and traits of several freshwater taxa, including larval amphibians (e.g., behavior, color, fin height, and length of intestine). Here, I tested the influence of pond canopy cover, floating vegetation, and pond morphology on the phylogenetic structure of an anuran metacommunity in the Atlantic Forest of Southeastern Brazil. I sampled tadpoles in 13 ponds and marshes from June 2008 and July 2009 in the Serra da Bocaina National Park, São Paulo. After building a metacommunity phylogeny, I used an eigenvector-based technique to describe the metacommunity phylogenetic composition (Principal Coordinates of Phylogenetic Structure, PCPS). I then run a db-RDA to evalute whether a subset of these eigenvectors can be explained by environmental variables. I found that pond canopy cover and floating vegetation were the main variables influencing lineage sorting in this metacommunity. Canopy cover separated hylid lineages from other families that were associated with open areas. Floating vegetation separated two hylid tribes (Cophomantini and Dendropsophini). Our results mainly suggest that the effect of canopy cover and floating vegetation on the structure of anuran metacommunity may affect not only species, but also entire lineages.

## Introduction

Phylogenetic information has been increasingly included into studies of (meta)communities (Webb et al. 2002, Mouquet et al. 2011, HilleRisLambers et al. 2012). The main argument for using phylogenies to infer community assembly mechanisms is that it would be a multivariate proxy for species’ traits, provided that traits that influence community assembly exhibit phylogenetic signal (Webb et al. 2002, Kraft et al. 2007, but see Losos 2012). When fulfilled these assumptions, environmental filters were claimed as the main mechanism in the assembly of clustered communities, whereas negative, density-dependent mechanisms (mainly resource competition) lead to overdispersed communities. This idea became both very influential and controversial throughout the years, but reinvigorated the study of community assembly (Mouquet et al. 2011, Kraft & Ackerly 2014).

Recently, the original framework proposed by Webb et al. (2002) has been challenged (see Fox 2012) following the results of experiments (Narwani et al. 2013, Fritschie et al. 2014, Godoy et al. 2014) that incorporated the theory of species coexistence as advocated by Chesson (2000). These studies with annual plants and green algae consistently found that competitive hierarchy does not correlate with phylogenetic distance. Therefore, competitive outcome could not be predicted simply from the phylogenetic relationships of species in a community. As a consequence, the interpretation of commonly used phylogenetic structure metrics, such as Net Relatedness Index (NRI), in which overdispersed communities are the result of competition and clustered ones caused by environmental filtering has been now undermined.

Regardless of these limitations, testing how environmental filtering affects lineage sorting can still be informative to understand processes governing community assembly (see Kraft et al. 2015). For example, it is still necessary to document the distribution of lineages throughout the landscape in response to environmental gradients, as a first evidence for environmental filtering (Kraft et al. 2015). Such studies are even more relevant when conducted with organism and ecosystems usually not evaluated under this perspective, such as pond-dweeling amphibians in tropical regions, as has been claimed elsewhere (Vamosi et al. 2009, Logue et al. 2011, Hortal et al. 2014). Such explorations would broaden our understanding of the many mechanisms acting on community assembly of a wider variety of organisms and help solve contigencies in the field.

Pond-dwelling amphibians are good model organisms to investigate metacommunity processes in freshwater ecosystems (Wilbur 1997), since they are subjected to different enviromental and spatial factors throughout their aquatic and terrestrial life stages (reviewed in Alford 1999, Wells 2007), which usually are scale dependent (e.g., Van Burskirk 2005). The dispersal is restricted to the adult stage, while the spatial distribution of larvae is determined by oviposition site selection by adults (Weels 2007). Furthermore, they inhabit discrete habitats, whose boundaries are easily identifiable, so there is little doubt about community limits and size (Blaustein & Schwartz 2001, De Meester et al. 2005).

Environmental gradients are key to determine amphibian species distribution at the landscape scale (Werner et al. 2007, Provete et al. 2014), especially hydroperiod (Wellborn et al. 1996) and pond canopy cover (Schiesari 2006, Werner et al. 2007). While adults are influenced by variables around ponds, such as canopy cover and inter-pond distance, which determine oviposition site selection (reviewed in Wells 2007), tadpoles are usually influenced by pond morphology variables (e.g., volume and aquatic vegetation). Pond canopy cover can control energy input to pond ecossytems, in which closed-canopy ponds are limited mostly to allocthonous production from the terrestrial environment. This type of pond may represent harsh environments for tadpoles as they usually have food resource of low quality, low light incidence, and consequent reduced algae productivity, and low oxygen concentration (Schiesari 2006). As a consequence, only a subset of species is able to oviposit in shaded ponds (Werner et al. 2007, Van Buskirk 2011).

However, little is known about the influence of pond canopy cover on lineage sorting of freshwater organisms and other facets of biodiversity, such as functional and phylogenetic diversities (see Hortal et al. 2014). Several key questions remain open, such as the interplay between these environmental filters and metacommunity processes to lineage sorting of pond-dweeling amphibians. Here, I tested how anuran lineages are sorted along the gradients of canopy cover and pond morphology in a metacommunity in Southeastern Brazil using an eigenvector-based technique to describe metacommunity phylogenetic structure.

## Material and methods

### Study site and sampling

This study was conducted on the highland portion of the Serra da Bocaina National Park, São José do Barreiro, São Paulo, southeastern Brazil (22° 40’ to 23° 20’ S; 44° 24’ to 44° 54’ W; Garey et al., 2014). Tadpoles were sampled in 13 ponds and marshes spread across 11 Km^2^ using a hand dipnet along the entire margins of water bodies, with effort proportional to surface area (Skelly & Richardson, 2010). Samplings occurred between June 2008 and July 2009. Ponds were separated by a maximum 7 Km (Fig. 1). Tadpoles were fixed in the field with 10% formalin. Ponds were sampled within a 1-week period each month.

**Fig. 1.**
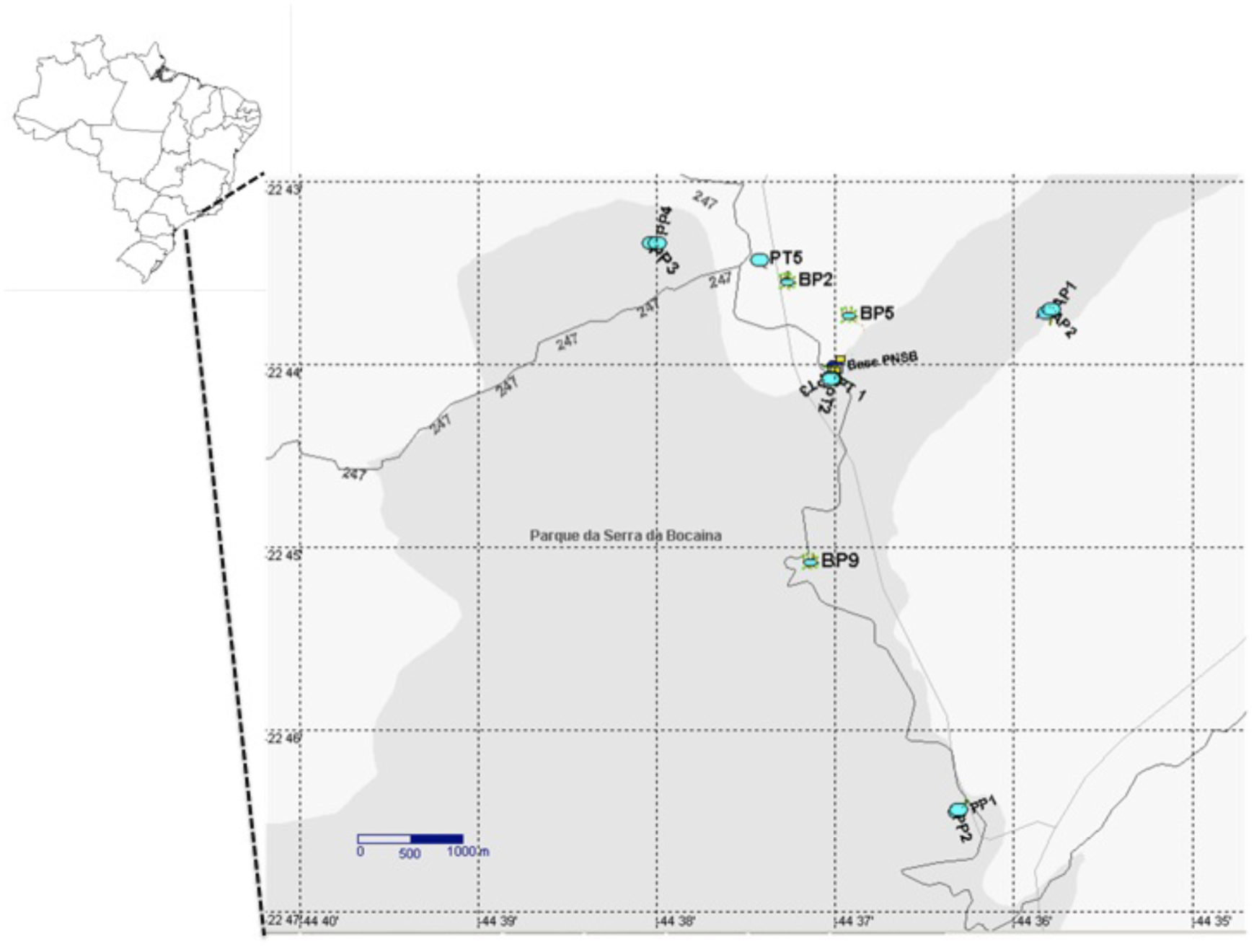
Spatial distribution of ponds sampled during this study in the Serra da Bocaina National Park.

I recorded variables of pond morphology during the rainy season: surface area (m^2^), maximum depth (m), besides floating aquatic vegetation (%), and pond canopy cover (%). I estimated it visually by dividing pond surface into quadrants. Pond canopy cover was measured using a spherical densitometer (Forestry Suppliers, Jackson, MS, USA); measurements were taken in four directions (N, S, E, W), and the center of the pond. A previous work (Provete et al. 2014) has found that water chemistry did not influence tadpole species abundance and composition at the landscape scale in this study site. Furthermore, water chemistry variables did not vary considerably at the landscape scale. Hydroperiod is an important environmental gradient in freshwater habitats (Wellborn et al. 1996). However, only 3 out of the 13 ponds were temporary (Provete et al. 2014). Therefore, these variables were not included in the further analyses. Prior to the analyses, I standardized environmental variables to zero mean and unit variance using the package vegan (Oksanen et al., 2013) in R version 3.1.2 (R Core Team, 2014).

### Metacommunity Phylogeny

The metacommunity phylogeny (Fig. 2) was created by prunning the dated tree of Pyron & Wiens (2013) for amphibians to include only the species found in the metacommunity using the function prune.sample of the R package picante (Kembel et al. 2010). That amphibian phylogeny was built under Maximum Likelihood criterion using the supermatrix approach from nucleotide sequences of nine nuclear and four mitochondrial genes available at GenBank. Two of those mitochondrial genes were shared by 90% of species, but there was considerable amount of missing data. Authors did not make available the whole set of the most likely trees, so it is not possible to promptly include phylogenetic uncertainty in the analysis. However, most of the relationships among deep nodes (e.g., representing splits among families; Duarte et al. 2016; Capítulo 2) are well supported and similar to previous topologies (e.g., Roelants et al. 2011). Therefore, there would be little difference among the most likely trees. I only used the first two PCPS in further analysis (see below). These PCPS capture variation deep in the tree and would be little affected if I incorporate different topologies. Disagreement among topologies are more common at the species and genera level, which are captured by PCPS with lower eigenvalue and not used in this study.

**Fig. 2.**
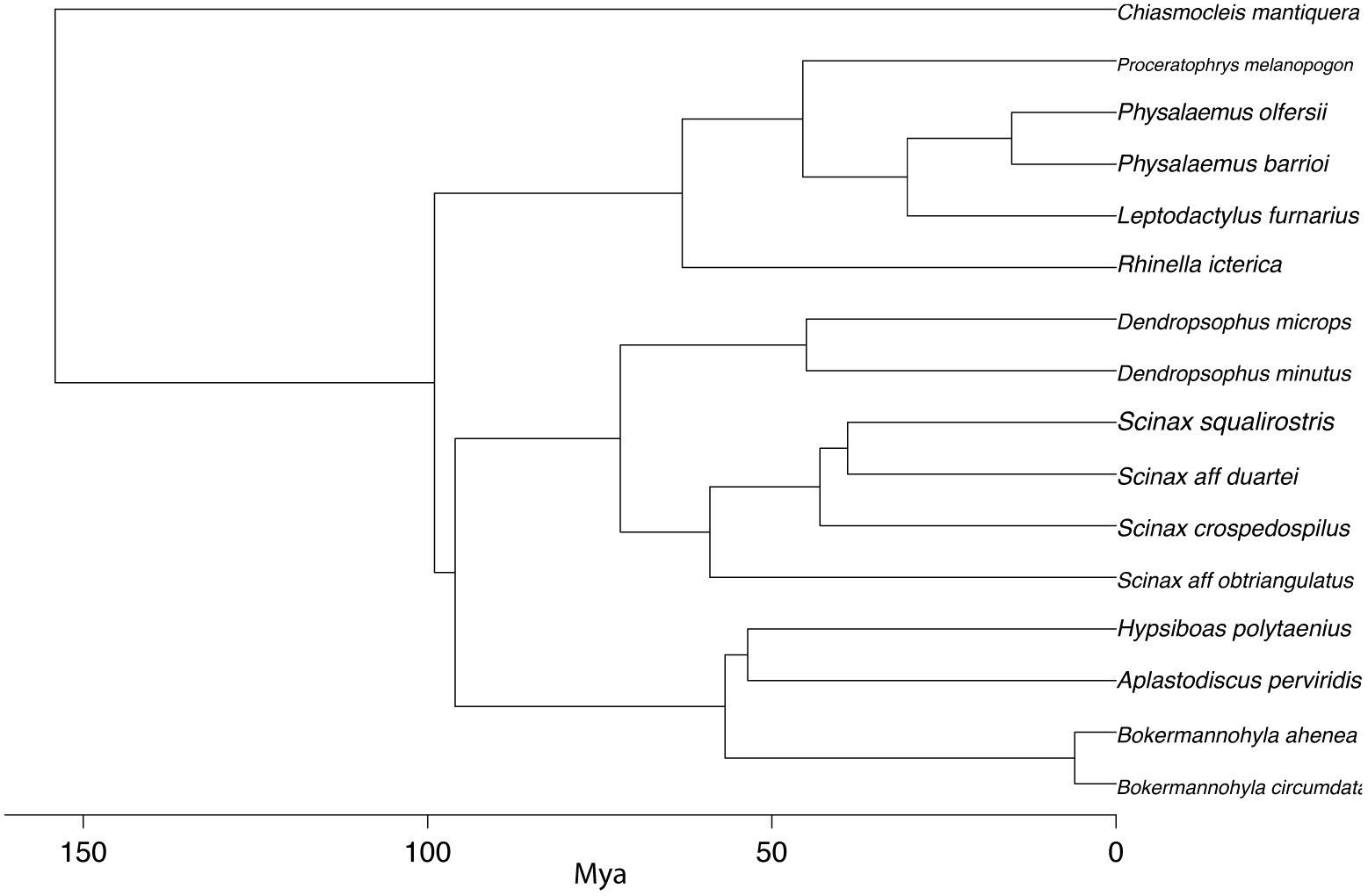
Metacommunity phylogeny for the study site at the Serra da Bocaina National Park pruned from the dated tree of Pyron & Wiens (2013).

I then calculated a patristic distance matrix (**D**) of the phylogenetic tree. The distances were converted into similarities following the formula: S_ij_=1-[d_ij_/max(d_ij_)]. Each element in matrix **S** was then divided by its column totals to produce a matrix **Q**. This matrix contains the phylogenetic weights that represent the phylogenetic belonging of each taxon to all others in the phylogeny (Pillar & Duarte 2010, Duarte et al. 2016). Then, the matrix **Q** was multiplied by the matrix **W** containing the site by species composition in the metacommunity to produce the matrix **P**, which contains the phylogeny-weighted species composition. Finally, a Principal Coordinates Analysis (Lengendre & Legendre 2012) was performed in the **P** matrix using squared-root Bray-Curtis dissimilarity. A PCoA calculated in the **P** matrix produces eigenvectors called Principal Coordinates of Phylogenetic Structure (PCPS, Duarte 2011, Duarte et al. 2016). These eigenvectors describe ortogonal phylogenetic gradients of the metacommunity and can be further used as response variables in multivariate analyses. Calculations were conducted in the R package PCPS (Debastiani & Duarte 2014).

### Data analyses

The procedure described above produced 12 PCPS (number of sites-1), with the first PCPS accounting for 83% of the variation in the matrix **P**. Since I was interested in understanding how lineages were affected by environmental variables, I run a distance-based Redundancy Analysis (db-RDA; Legendre & Legendre 2012) using the eigenvectors as predictors. However, two nearby sites may have more similar environmental variables than two far apart sites, i.e., they may exhibit spatial correlation (Legendre 1993). Consequently, I included spatial variables in the form of Moran Eigenvector Maps (MEMs; Legendre & Legendre 2012) to account for the spatial arrangement of sites based on a Delaunay triangulation. Then, I calculated the neighborhood matrx and converted it into a spatial weighting matrix, with each edge having weight depending on the linear distance between nodes. Lastly, I calculated MEMs, which generated 12 eigenvectors of which 3 had significantly positive Moran’s *I*. Thus, I only included the first three MEMs in the further analysis.

Here, I followed Duarte et al. (2012) and Anderson & Willis (2003) recomendation to select PCPS to be entered in the analysis. Briefly, I run several db-RDA with the same environmental variables, but with an increasing number of PCPS and chose the subset with the highest F value, indicating the best fit (Duarte et al. 2012). This procedure maximizes the association between eigenvectors and the predictor variables. The first two PCPS were selected using this procedure. Analysis was conducted in the R package vegan (Oksanen et al. 2013) using the function capscale. I also implemented a stepwise selection of environmental variables using the ordistep function of the vegan package. Then I use variation partitioning to disentangle the relative contributions of the reduced environmental and spatial model in predicting the phylogenetic structure of the metacommunity, as captured by the first 2 PCPS.

## Results

I collected about 29,000 tadpoles of 16 anuran species, belonging to five families. Species distribution suggests that most species are canopy generalists, whereas only two are specialists, *Chiasmocleis mantiqueira* and *Physalaemus olfersii* (Fig. 3). Indeed, the final best model included canopy cover and floating vegetation, besides the three MEMs (Fig. 4). The pure environmental component explained 53% (*P*=0.033), while the pure spatial component explained 45% (*P*=0.029) of the variation in the metacommunity phylogenetic structure (Fig. 5), with the spatiall structured environmental variation having an insignificant contribution.

**Fig. 3.**
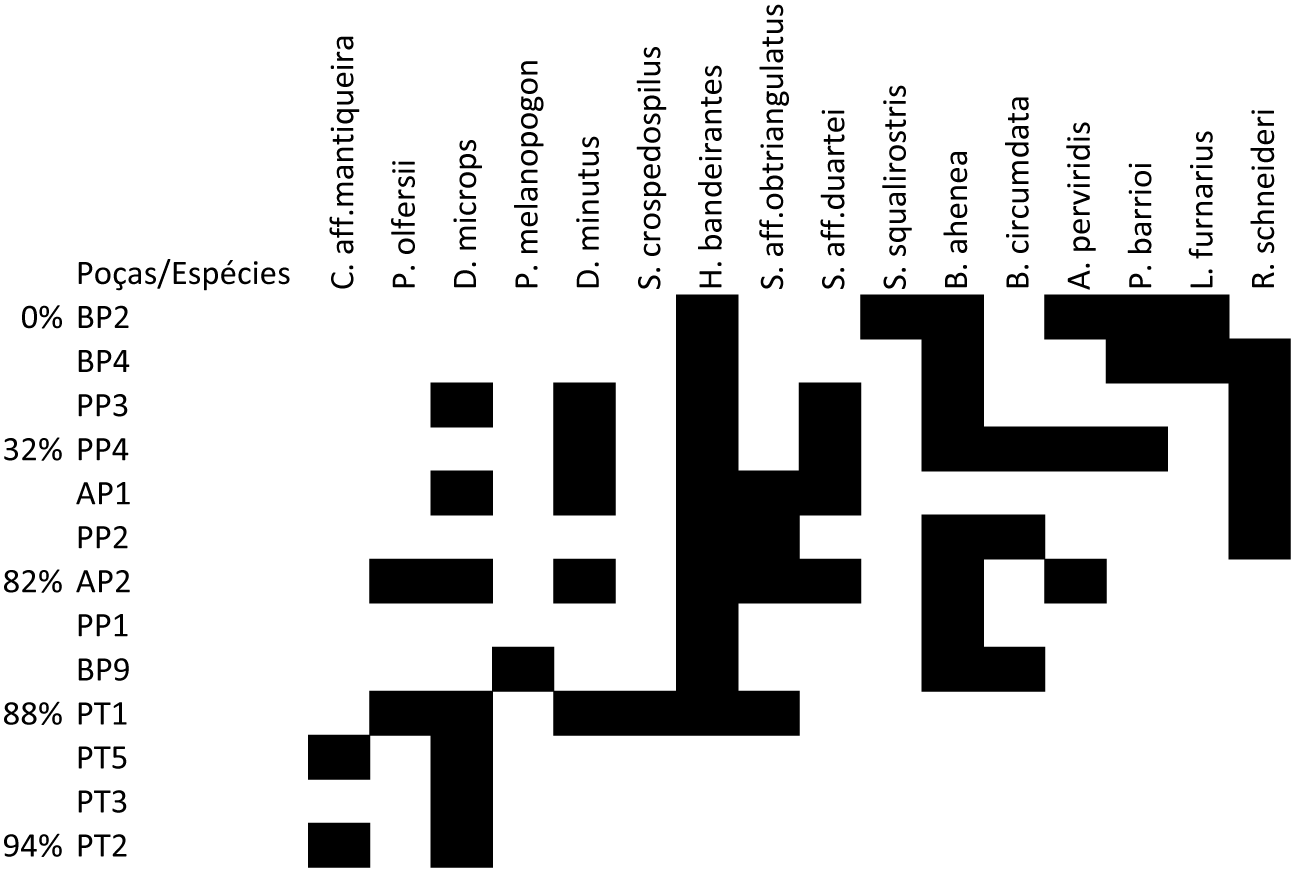
Species distribution in relation to canopy cover. Ponds are arranged from low canopy cover (on the top) to high canopy cover (bottom). Percentages refer to canopy cover. See Fig. 2 for full species names.

**Fig. 4.**
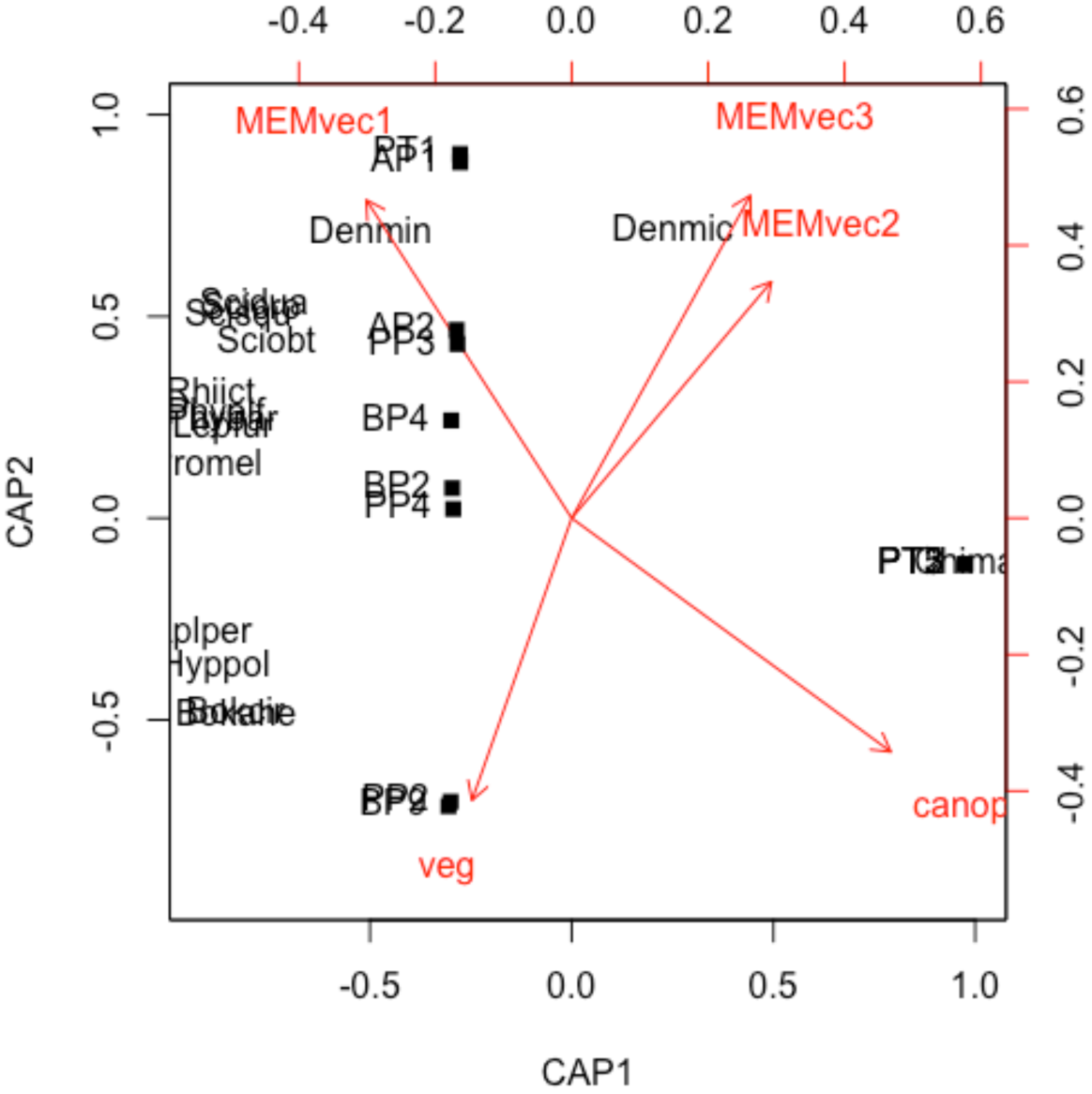
Ordination diagram of the first two axis of the db-RDA, showing the main variables affecting the metacommunity phylogenetic structure (arrows). Species names are abbreviated, see Fig. 2 for full species names. Squares represent ponds sampled.

**Fig. 5.**
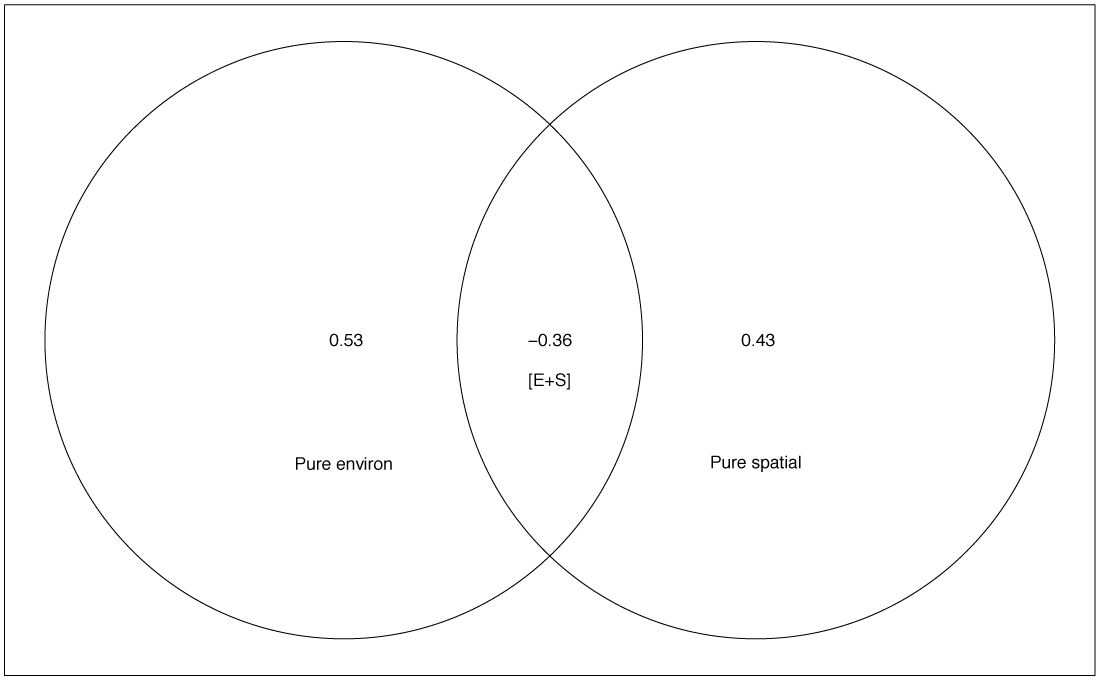
Result of variation partitioning of the Principal Coordinates of Phylogenetic Structure between the environmental (canopy and floating vegetation) and spatial (first three MEMs).

The tribe Cophomantini of hylids were associated with increasing pond canopy cover and floating vegetation, while Dendropsophini and the remainig clades (Leptodactylidae, Odontophrynidae, and Bufonidae) were associated with open areas (Canonical R^2^ _adj_=0.596; F_5,7_=4.555; *P*=0.029; Fig. 4).

## Discussion

Canopy cover seems to be a strong filter not only for species (e.g., Werner et al. 2007, Provete et al. 2014), but also for entire clades (this study). Furthermore, I found that its influence occurs at deep levels of the phylogeny and that arboreal species (hylids) are differently affected by canopy cover. These results agree with previous studies on both temperate (Schiesari 2006, Werner et al. 2007, Van Buskirk 2011) and tropical ecosystems (e.g., Provete et al. 2014) that found that only a small subset of species are able to occupy shaded ponds, but those that do might benefit from low exploitative competition.

Differences in habitat use among clades maybe the main mechanism driving this pattern, but also traits related to reproductive strategy and density-dependent interactions at the larval stage. For example, most ground-dwelling species from the families Leptodactylidae, Odontophrynidae, and Bufonidae occurred in open areas. These species usually take advantage of the flush of primary productivity available in open canopy ponds (Wassersug 1975, Wilbur 1980, 1997), so tadpoles can reach metamorphosis rapidly before the pond dries. They also have reproductive modes adapted to withstand direct sunlinght exposure, such as foam nests (Wells 2007, Crump 2015). Conversely, forest-dwelling species have traits that allow them to explore the vertical stratum of the forest, such as adhesive discs. Both adhesive discs (Wells 2007) and foam nests exhibit a strong phylogenetic signal (Gomez-Maestre et al. 2012), each occurring in only one family of this community phylogeny, Hylidae and Leptodactylidae respectively. As a result, conserved adult traits in response to environmental gradients, which in turn influence breeding site selection, seem to ultimately determine tadpole spatial distribution (Provete et al. 2014, this study).

Interestingly, there seems to be some generalist (e.g., *Dendropsophus microps*) but only a few specialist species (e.g., *Chimasmocleis mantiqueira*) regarding the occurrence along the pond canopy cover gradient. In the studied site, *C. mantiqueira* was an explosive breeders and very abundant in temporary, closed canopy ponds formed during the rainy season. This species belongs to an early-split in the phylogeny, separating two superfamilies. The first eigenvectors used capture variation near the root of the phylogeny (Capítulo 2) and could also be influencing the separation of this species from the other clades. Conversely, *L. furnarius* and *R. icterica* were restrictied to open-canopy ponds. Several lines of evidence have suggested that not only species composition, but also behavior, color, morphology, and overall performance of anuran larvae vary along the canopy cover gradient (Schiesari 2006, Werner et al. 2007, Van Buskirk 2011). These seem to be the result of varying resource quality and quantity (Schiesari 2006) along the canopy cover gradient. As the results of this study suggest, these same factors could be at play in tropical enviroments, but further field experiments are needed to test this hypothesis.

My results reinforce the findings of a previous study in the same site (e.g., Provete et al. 2014), which report the spatial dependence of canopy cover and floating vegetation at a broad scale as main determinants of tadpole metacommunity structure. This study analysed taxonomic composition as a measure of diversity and included environmental variables as proxy for adult breeding site selection. Thus, it seems that those variables may affect both phylogenetic and taxonomic composition. This raises the question about what traits drive this pattern (see Both et al. 2011 for an example with pond hydroperiod).

Further research investigating the role of phylogeny on amphibian community assembly should include adult traits related to habitat use and reproductive modes as well as larval morphology. It is likely that reproductive modes of anurans (Wells 2007, Crump 2015) play a key role determining tadpole spatial distribution at the landscape scale, since they seem to be a very conserved trait at the clade level (Gomez-Mestre et al. 2012). In addition to affecting the performance of larvae (Williams et al. 2007), it is known that morphological traits of larval anurans vary along the canopy cover gradient (Van Buskirk 2011, Stoler & Relyea 2013). This environmentally driven phenotypic plasticity related to the occupation of shaded ponds suggests that some larval traits are likely not constrained by the phylogeny. Thus, apparently the environment also plays a role in driving community assembly of larval anurans along this gradient. But formal tests of coexistence (Chesson 2000, HilleRisLambers et al. 2012), i.e., estimating species’ vital rates and competition coeficients would help determining if these plastic traits are correlated either with stabilizing niche differences or equalizing mechanisms, i.e., conferring a competitive advantage for species that are more plastic.

## Acknowledgements

This work was partially funded by a master’s fellowship from the Fundação de Amparo à Pesquisa do Estado de São Paulo (Proc. 08/55744-6) granted to DBP. Michel V. Garey helped in the fieldwork. IBAMA provided collecting permits (#14474) and the staff of the Serra da Bocaina National Park provided logistical support and housing. DBP received a PhD fellowship from CAPES-DS.

